# Real-time Bayesian optimization of deep brain stimulation for personalized cognitive control enhancement

**DOI:** 10.64898/2025.12.30.697057

**Authors:** Evan M. Dastin-van Rijn, Elizabeth M. Sachse, Michelle Buccini, Bradley Angstadt, Jonathan Bennek, Megan E. Mensinger, Alik S. Widge

## Abstract

**Background:** Identifying effective deep brain stimulation (DBS) parameters for psychiatric disorders has historically been a time-consuming and error prone process due to a lack of an objective and rapid readout of target circuit engagement. Cognitive control may have use as a biomarker of treatment efficacy but it has yet to be shown that DBS parameters can be reliably optimized to produce cognitive control improvements in individual subjects.

**Objective:** We sought to leverage a rat model of DBS-driven cognitive control improvements to determine whether state of the art optimization algorithms could consistently identify effective stimulation amplitudes to enhance cognition.

**Methods:** We delivered periods of active and inactive DBS-like stimulation at variable parameters while rats performed a Set-Shifting task that we previously showed to be stimulation-sensitive. We tested both predefined settings of interest and settings that were personalized to individual animals using Bayesian Optimization. Measurements of task performance including reaction time and accuracy were compared between acute, optimized, and traditional settings to evaluate effects on cognitive control.

**Results:** Acute stimulation reduced reaction times without hindering accuracy (N=15), replicating the effects previously observed with chronic stimulation. In a second cohort (N=6), optimization of stimulation amplitude successfully reduced reaction times in all animals with comparable effect size to historically best settings.

**Conclusion:** These findings confirm that optimization techniques can be effective for improving symptomatically-relevant cognitive markers supporting the feasibility of personalized, quantitatively-informed approaches to neuromodulation and target engagement for psychiatric and/or cognitive disorders.

**Highlights:** Proving target engagement is a substantial challenge across brain stimulation modalities, and objective, rapid behavioral read-outs may be a solution to that challenge.

Reaction times in cognitive control tasks are an example of a behavioral measure that changes rapidly in response to changes in stimulation parameters, and that also may predict clinical outcomes.

Individually optimal stimulation amplitudes for reducing reaction time by stimulating corticofugal fibers passing through the striatum can be determined using Bayesian Optimization.

Individually optimized settings discovered in an acute preparation demonstrate consistent effects when applied chronically.

## Introduction

At least 20% of patients with mental illnesses find no relief from standard care [1]. For those patients, neuromodulation therapies are a promising treatment option with the potential for precision and personalization – but that potential has not yet been realized [2]. Deep brain stimulators (DBS) are programmed on the basis of infrequent, subjective evaluations of mood while non-invasive alternatives are often targeted based on skull landmarks rather than brain anatomy, and use energy patterns that may not produce reliable physiologic effects [3–5]. This leads to a problem of target engagement – many neuromodulation therapies may fail for individual patients because the wrong pattern or intensity of energy was delivered, and thus the target brain circuit did not change. In motor disorders, where DBS has robust efficacy, stimulation has immediate and quantifiable effects on behavior [6]. Those immediate effects allow reliable target engagement, enabling both symptom relief and biomarker discovery that in turn drove novel closed-loop therapies [7,8]. In contrast, neurostimulation for psychiatric indications has been substantially hindered by a lack of similar target engagement markers [2,9].

We have proposed that target engagement could be verified, and physiologic markers discovered, by considering neurostimulation’s effects on cognitive control [2,10,11]. Both invasive modalities such as DBS and non-invasive modalities such as transcranial magnetic stimulation (TMS) can improve cognitive control task performance, in both diagnosed patients and healthy volunteers [12,13]. Cognitive control describes an individual’s ability to withhold a prepotent response in favor of a more adaptive, goal-oriented response. Deficits in cognitive control are characteristic of many psychiatric disorders and these deficits have been shown to diminish as symptoms improve [14–16]. Therefore, enhancing cognitive control could be an effective treatment target with a clear relationship to disorder symptomatology [17,18]. Additionally, unlike mood, cognitive control can be rapidly and objectively assessed via behavioral tasks [19].

Showing a group-level effect of stimulation on cognitive control does not automatically mean that we can produce that effect for individuals. For example, for internal capsule stimulation(which improved cognitive control in [12,17,20], anatomic heterogeneity means that no single electrode placement or set of parameters will reliably engage the target circuit in all patients [21]. That heterogeneity might be overcome, however, by using Bayesian optimization to identify patient-specific parameters [18,22].

Bayesian methods have shown promise in pain [23], motor [24–26], epilepsy [27], and neuroprosthetic [28,29] settings. These algorithms explore parameter space in an information-seeking fashion, building a map of a patient’s individual response to stimulation. By pairing such an optimizer with real-time monitoring of cognitive task performance, a clinician could identify stimulation settings that robustly enhance a circuit engagement biomarker such as performance on a cognitive control task.

We previously conducted simulations with noise patterns and effect sizes derived from human data, suggesting that optimizing DBS to enhance cognitive control is at least theoretically possible [18]. However, these simulations were limited to a single parameter (stimulation site) and relied on assumptions about trial-to-trial correlations and transient stimulation effects. We have also seen acute effects of stimulation on cognitive control in humans but the exact timing of these effects, how they might translate between patients/applications, and their impact on optimization performance is uncertain [17]. While there is substantial evidence supporting the feasibility of stimulation optimization to enhance cognitive control, these ideas have never been tested *in vivo*.

To close that gap we leveraged our existing rat model of cognitive control enhancement using DBS-like electrical stimulation. We confirmed that the enhancement we previously observed could be replicated acutely, enabling rapid assessment of the effect of DBS-like stimulation on cognitive control. As in humans, these acute effects were sensitive to the choice of stimulation parameters, confirming that differences between effective and ineffective parameters could be measured acutely. Based on these two findings, we ran a proof of concept Bayesian optimization experiment where the ideal stimulation amplitude for enhancing cognitive control was identified for each rat. Optimization consistently and robustly converged to effective parameters. When parameters identified through acute optimization were applied continuously, they replicated our previously reported behavioral effect. These findings confirm that black-box techniques can be effective for improving symptomatically-relevant cognitive markers, even for relatively noisy markers such as trial-level response times. Collectively, these results support the feasibility of personalized approaches to neuromodulation for psychiatric and/or cognitive disorders.

## Methods

### Animals

12 male Long-Evans rats (250-300 g) and 14 Long-Evans female rats (200-250 g) were obtained from Charles River Laboratories. Based on past experiments [20,30], we targeted n=8 for each analysis with the goal of having at least 6 rats per group based on expected effect sizes. All experiments were approved by the University of Minnesota Institutional Animal Care and Use Committee (protocols 1806-35990A and 2104-39021A) and complied with National Institutes of Health guidelines.

### Set-Shifting task

The operant Set-Shift task (Fig. S1A) was adapted from [31] and described previously [20]. Briefly, rats alternated between two rules: a cue-based “light” rule and a spatial “side” rule. Each trial began when the rat poked the illuminated center port, after which one peripheral port was lit. Under the light rule, rats responded to the illuminated port regardless of location; under the side rule, they responded to a fixed spatial location regardless of illumination. Correct responses were rewarded with one pellet. After five consecutive correct choices, the rule switched to the opposite dimension. Shaping and food deprivation procedures were as in [20,30].

Unlike earlier versions, we fixed the trial count (150–160 trials per session) rather than number of shifts. This design, also used in the mouse adaptation [32], facilitated consistent analyses. Stimulus presentation and sequencing were controlled by a *pybehave*/OSCAR [33,34] framework operating Coulbourn Instruments chambers.

### Surgery

Rats were anesthetized with isoflurane and mounted in a stereotaxic frame. We exposed the skull and drilled a burr hole to implant a bipolar stimulating electrode targeting the mid-striatum (+1.4 AP, ±2.0 ML, -5.8 DV) [20,30].

### Histology

At the end of the experiment, rats were sacrificed for histology. Electrodes were localized using a fluorescence microscope (Keyence, Osaka, Japan). Images of tracts were overlaid on a rat brain atlas to determine the approximate coordinates (Figure 1A). Electrodes were considered on-target when the tips of the electrodes were within the mid-striatum region defined by previous work [20].

**Figure 1:**
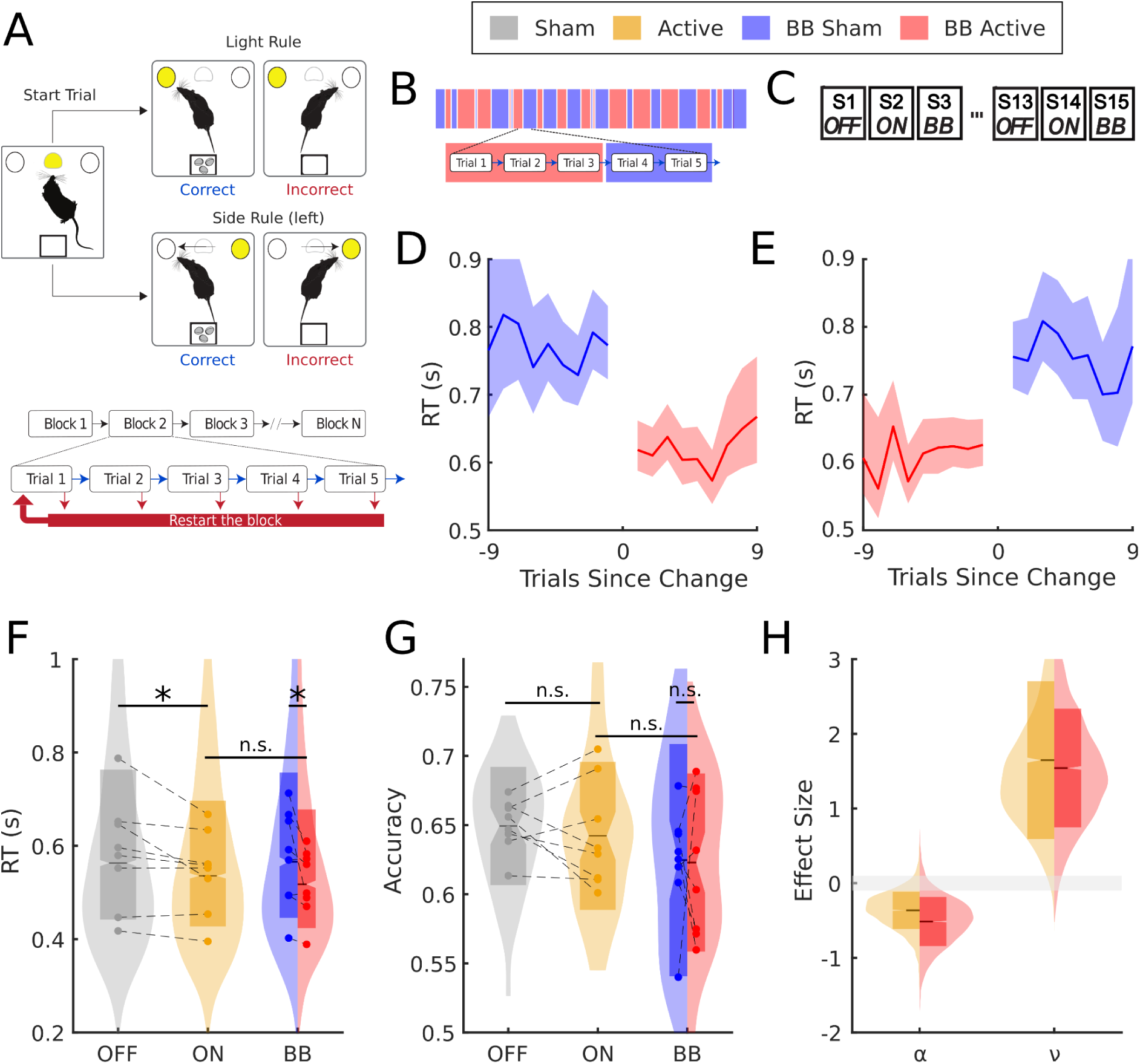
Trial-based stimulation replicates the cognitive control enhancement from whole-task stimulation. **(A)** The Set-Shift task. Rats initiate a trial by poking the middle port and then must select either an illuminated port (light rule) or ignore the light and select a specific side of the chamber (side rule). The rules are not cued, and shift to a new rule after the rat has sequentially completed 5 correct trials of the current rule. Errors reset to the beginning of the current 5-trial block. **(B)** Illustration of block-based sequencing in the set-shift task. Set-shift trials were grouped into blocks of 1, 3, 5, 7, or 9 trials. For the duration of each block, stimulation was either continuously active (red) or continuously off (blue). Blocks alternated between on and off conditions over the duration of the task. **(C)** Illustration of the sequence of task conditions for the block-based experiment. Rats completed 5 sessions each of continuous off (OFF), continuous on (ON, 300 μA, 50 μs, 130 Hz), or block-based (BB) stimulation for a total of 15 sessions. Sessions were completed in a predefined repeating sequence of OFF/ON/BB. **(D & E)** Change in reaction time (RT) relative to changes in stimulation condition during the block-based task.**(D)** Following a change in settings, active stimulation (red) produced an immediate and lasting reduction in reaction time compared to inactive stimulation (blue). **(E)** Similarly, a change from active to inactive stimulation immediately reversed this decrease. **(F)** Distributions of reaction time as a function of task condition. Mean values for individual rats in each condition are shown by the scattered points and dashed lines link points from the same rats between conditions. Distributions were computed over all trials. **(G)** Distributions of accuracy as a function of task condition. Distributions were computed over all sessions. **(H)** Distributions of stimulation effect on RLDDM boundary separation and drift rate over 4000 posterior draws. The median of the distribution is indicated by the solid line. The gray shaded area around zero represents the ROPE for a null effect (effect size less than 0.1). *P < 0.05; all P values represent Wald Z tests of model parameters from GLMMs. See supplemental tables for formulae and statistical details.

### DBS-like electrical stimulation

Electrical stimulation was delivered using a StimJim dual-channel electrical stimulator ([35], assembled by Labmaker, Berlin, Germany) controlled by py*behave* [33]. Stimulation used bipolar, biphasic, charge-balanced square wave pulses ranging from 0-350 μA, at 20 or 130 Hz, with a pulse width of 50 µs per phase, totaling 100 µs per pulse. 130 Hz is a typical human DBS frequency [3] and was used in previous experiments in both humans [12,17] and rats [20,30] while 20 Hz stimulation typically has reduced effects in motor studies [36]. Stimulation parameters varied during and across sessions depending on the task condition (see below). We confirmed that stimulation waveforms were balanced and consistent using an oscilloscope and 1 kOhm series shunt resistor.

### Block-based stimulation

In our original experiments, stimulation was either continuously on or off [20]. To assess acute effects, we developed a block-based protocol. Each session was divided into contiguous blocks of 1, 3, 5, 7, or 9 trials, independent of task rules. Alternating blocks were assigned active or inactive stimulation, with at least three instances of each active/sham block length per session. Block sequences ensured equal transitions between block lengths, and all rats experienced identical sequences for corresponding Set-Shift protocols. Stimulation parameters were updated between blocks, 3.5 s into the inter-trial interval.

To compare acute and whole-task effects, we interleaved continuous-stimulation with block-based sessions. Each condition (OFF, ON, Block-Based) was tested across five sessions per rat in pseudorandom order (e.g., OFF1, ON2, BB3, OFF4…).

### Variable parameter stimulation

To determine whether acute effects from block-based stimulation depended on specific parameters, we conducted a follow-up experiment varying stimulation amplitude and frequency within sessions. Five conditions were tested: stimulation off, 100 µA at 130 Hz, 200 µA at 130 Hz, 300 µA at 130 Hz, and 300 µA at 20 Hz. Each Set-Shift session was divided into contiguous blocks of 1, 3, 5, or 7 trials, with adjacent blocks assigned different conditions. Every session included two instances of each block length per condition. To ensure adequate power for model fitting, five sessions of variable-parameter stimulation were run per rat, using each Set-Shift protocol once.

### Statistical analysis of Set-Shift behavior

All analyses were performed in MATLAB (R2021b). Generalized linear mixed-effects models (GLMs) were used to analyze Set-Shift behavior, as they accommodate repeated measures, non-Gaussian response and binary data, and maintain comparability with prior work [12,17,20]. GLMs included fixed effects for stimulation condition (OFF or categorical acute parameters), task condition (OFF, ON, or BB), and rule type (side or light), with random effects for rat, session, and protocol. Omission trials—where no response occurred within 3 s—were excluded except when omissions were analyzed directly. Full GLM specifications are provided in the Supplemental Tables. Statistical significance was defined as *p* < 0.0083 (Bonferroni correction: 0.05/6 task measures). For the optimization follow-up, only reaction time and accuracy were tested (correction factor = 2).

### Reinforcement Learning-Drift Diffusion Model of Set-Shift Behavior

To confirm that acute stimulation reproduced the effects of whole-task stimulation, we adapted a reinforcement learning–drift diffusion model (RLDDM) [20,37,38]. This model integrates a drift-diffusion process, capturing response times and choices via latent decision variables, with reinforcement learning that updates values based on reward history. On each trial, the RLDDM assigns learned values to left and right nose pokes according to location and illumination. The value difference determines the drift rate and bias, while boundary separation specifies the evidence required for a decision. After each choice, side and light values are updated through reinforcement learning (see Figure 3A in [20]).

Based on prior results, we modeled stimulation effects on drift rate, boundary separation, bias, and non-decision time only. After validating model fit via posterior predictive checks, we quantified stimulation effects using the region of practical equivalence (ROPE), probability of direction, and Cohen’s *d* [39].

### Bayesian optimization of Set-Shift reaction times

In a cohort of seven rats, we conducted a proof-of-concept study to identify the stimulation amplitude minimizing Set-Shift reaction times using Bayesian optimization. This method sequentially samples the parameter space at informative locations, selecting each new sample based on predictions from a surrogate model and their associated uncertainty [40,41]. Optimization ran in parallel with the Set-Shift task for five days per rat (155 trials per session, 775 total). After each trial, the log reaction time and stimulation amplitude were added to prior data to fit a Gaussian process (GP). The GP informed an acquisition function balancing exploration and exploitation, and the next amplitude was chosen by maximizing this function. Parameters were updated 3 s into the inter-trial interval, with computations typically requiring <1 s (Intel i9-12900K, 64 GB RAM, NVIDIA RTX 4070 12 GB).

As a control, five sessions interleaved between optimization days used a space-filling Sobol sequence for amplitude selection [42]. After optimization, we compared each rat’s individualized optimal setting with stimulation-off and standard 300 µA conditions across five pseudorandomized sessions per condition to balance Set-Shift protocols (e.g., OFF1, ON2, OPT3, OFF4…).

Based on prior simulations [22], we used a hybrid approach combining the GIBBON acquisition function [43] with an ε-greedy mean/variance heuristic [44]. Because reaction times are noisy and require many samples to resolve underlying trends, we used a boundary-avoiding iterated Brownian bridge kernel (IBBK) [22,45] with a custom sigmoidal mean function to ensure optima could be detected even at parameter boundaries.

### Statistical analysis of optimization performance

We tested whether optimization and space filling yielded similar estimates of the optimal stimulation amplitude using bootstrapped regressions. For each rat’s dataset, residuals were obtained by subtracting the GP estimate from the raw samples. Residuals were then resampled with replacement, added back to the GP estimate, and the GP re-fit. Each bootstrapped GP was minimized to estimate the optimal amplitude for both optimization and space-filling data. We generated 1000 bootstrap samples, each providing one amplitude per method across the seven rats. For each sample, we computed an Orthogonal Distance Regression (ODR) using *scipy* [46]. ODR was used instead of ordinary regression since both variables contained measurement error; expected errors were set to the standard deviations of amplitudes for each method. Confidence intervals were computed from the bootstrapped regression distributions.

To assess convergence, we re-fit GPs using progressively larger subsets (in steps of 50 trials) of each dataset to estimate model recommendations over the course of optimization. Each GP was minimized to obtain the amplitude that would have been proposed if optimization had stopped after each subset. For each rat, intermediate proposals were compared to the final model (fit to all data) to estimate absolute error relative to the converged solution. Variance in convergence over time was estimated using 9,999 bootstrapped samples drawn with replacement from mean convergence trajectories for each method.

### Assessing multivariate outcomes during space filling

Because samples during space filling are evenly distributed across parameter space, we fit Gaussian processes (GPs) for multiple behavioral outcomes—reaction time, accuracy, and initiation delay. For each rat and outcome, GPs with radial basis function kernels and fully optimized hyperparameters were fit.

Reaction and delay times were log-transformed and modeled using *botorch*’s *SingleTaskGP*, *ExactMarginalLogLikelihood*, and *fit_gpytorch_mll* [47]. Accuracy, being binary, was modeled with a Bernoulli likelihood using *SingleTaskVariationalGP* and a *VariationalELBO* objective [48]. Posterior means from the three outcomes were used to compute Pareto fronts illustrating trade-offs in performance as a function of stimulation amplitude.

## Results

### Effects of DBS-like stimulation on cognitive control can be measured acutely

Rats were implanted with stimulating electrodes in the right mid-striatum where we previously observed cognitive control enhancement from DBS-like electrical stimulation (Figure S1A-B) [20,30]. As in our prior work, we probed cognitive control using a Set-Shift task where rats must discriminate between cue (light) and spatial (side) rules (Figure 1A) [20,31]. Unlike prior versions of the task, we ran a variant where stimulation varied within a single session with trials being grouped into on and off blocks (Figure 1B). These block-based, acute sessions were compared to the continuous active/sham approach we used in previous studies (Figure 1C). Acute stimulation produced an immediate and lasting reduction in RT (Figure 1D) that was immediately reversed when stimulation was turned off (Figure 1E). This acute RT reduction was significant (*β* = −59 ms and *P* = 2.81 × 10^−14^) and comparable to the reduction seen with continuous stimulation (*β* = −61 ms and *P* = 1.56 × 10^−3^, Figure 1F, Table S1). As was the case for continuous stimulation (*P* > 0.05), this RT improvement did not come at the cost of any reduction in accuracy or rate of task progression (*P* > 0.05, Figure 1G & S2A, Table S2 & S3). Both acute and continuous stimulation also significantly reduced the time taken to initiate a trial (*P* < 2.15 × 10^−7^, Figure S2B, Table S4) and the percentage of trials with omissions (*P* < 5.46 × 10^−3^, Figure S2C, Table S5) as seen for continuous stimulation. We observed some evidence of side-biasing from the unilateral implant (*β* = 0.100 and *P* = 0.04), but neither acute nor continuous stimulation showed any significant stimulation-related change in side bias (*P* > 0.05, Figure S2D, Table S6).

We further confirmed that the RT reduction from acute, block-based stimulation evoked the same cognitive mechanism as continuous stimulation. We fit the same Reinforcement Learning-Drift Diffusion model (RLDDM) as our previous work [20,37]. Simulated responses from the RLDDM strongly matched empirical behavior and psychometrics for both acute and continuous data (Figures S3A-C). As in our prior cohort, DBS-like stimulation substantially lowered boundary separation [probability of direction (pd) = 94.73%, median = −0.338, and 10% in region of practical equivalence (ROPE)] and substantially increased drift rate (pd=99.2%, median=1.05, 1.46% in ROPE) (Figure 1H). Acute stimulation showed a comparable decrease in boundary separation (pd=94.43%, median=-0.495, 4.75% in ROPE) and increase in drift rate (pd=98.68%, median=1.46, 1.1% in ROPE) (Figure 1H). Neither continuous nor acute stimulation showed any substantial effect on bias or non-decision time (Figure S3D). This modeling suggests that the general improvement in conflict resolution from continuous stimulation is present even when stimulation is delivered acutely.

### Acute effects of DBS-like stimulation on cognitive control are sensitive to the choice of parameters

To confirm that the acute effects we observed were sensitive to the choice of stimulation parameters, we ran a variant of the block-based task in the same rats where stimulation amplitude and frequency were allowed to vary over blocks within each session (Figure 2A). Stimulation amplitudes of at least 200 μA were sufficient to significantly reduce RT and this reduction was present both for 130 Hz high frequency stimulation and 20 Hz low frequency stimulation (*β* < −49 ms and *P* < 1.52 × 10^−7^, Figure 2B, Table S7). This same pattern was present for initiation delay (*P* < 6.66 × 10^−23^, Figure S4A, Table S8) and omission percentage (*P* < 1.03 × 10^−3^, Figure S4B, Table S9). 200 μA stimulation at 130 Hz also significantly improved accuracy (*β* = 0.192, *P* = 0.031) while other settings showed no effects on accuracy (*P* > 0.05, Figure 2C, Table S10). As in the prior block-based experiment, we observed evidence of side-biasing from the unilateral implant (*β* = 0.108 and *P* = 0.05), but 300 μA stimulation at both frequencies reversed this side-biasing (*β* < -0.203 and *P* < 0.01, Figure S4C, Table S11). No setting had any significant effect on rule progress (*P* > 0.05, Figure S4D, Table S12). Lastly, we confirmed that the reduction in RT was independent of prior settings, i.e. that the RTs during a given stimulation setting were consistent regardless of parameters of the preceding block (Figure 2D, Figure S5).

**Figure 2:**
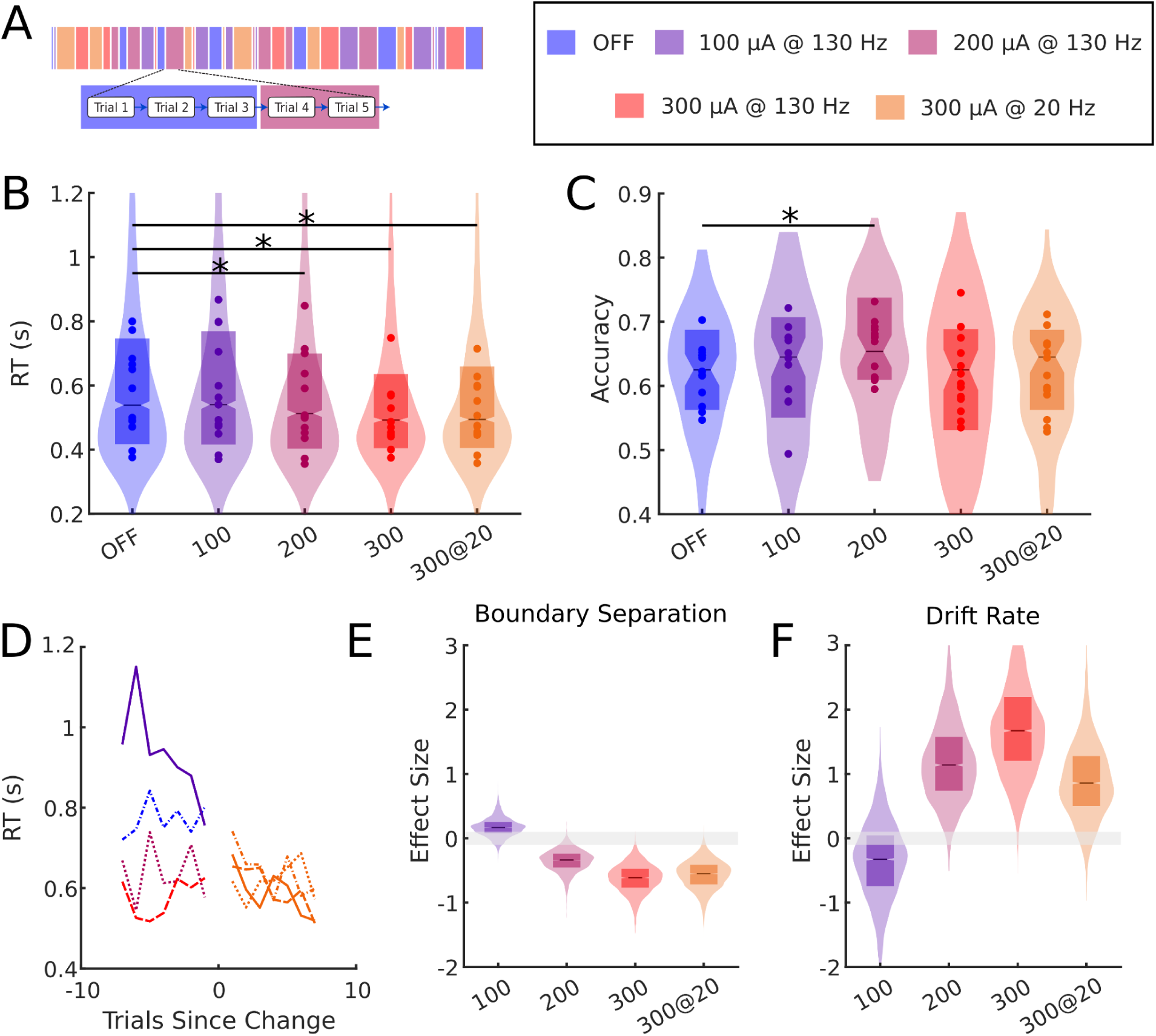
Efficacy of trial-based stimulation depends on stimulation dose. **(A)** Illustration of block-based sequencing with variable parameters in the set-shift task. Set-shift trials were grouped into blocks of 1, 3, 5, or 7 trials. For the duration of each block, stimulation was either continuously off (blue) or continuously on at a defined setting (purple=100 μA at 130 Hz, pink=200 μA at 130 Hz, red=300 μA at 130 Hz, and orange=300 μA at 20 Hz). Blocks switched between parameters over the duration of the task. **(B)** Distributions of reaction time as a function of stimulation parameters. Mean values for individual rats in each condition are shown by the scattered points. Distributions were computed over all trials. **(C)** Distributions of accuracy as a function of task condition. Distributions were computed over all sessions. **(D)** Change in reaction time (RT) when parameters were updated to the 300 μA at 20 Hz condition. Reaction times were averaged separately by the parameters that preceded the 300 μA at 20 Hz condition. Paired pre and post change averages are indicated by the line-style. **(E & F)** Distributions of stimulation effect on boundary separation (E) and drift rate **(F)** over 4000 posterior draws. The median of the distribution is indicated by the solid line. The gray shaded area around zero represents the ROPE for a null effect (effect size less than 0.1). *P < 0.05; all P values represent Wald Z tests of model parameters from GLMMs. See supplemental tables for formulae and statistical details.

To confirm that individual stimulation settings also engaged the cognitive mechanism identified previously, we fit the same RLDDM once again. Simulated responses from the RLDDM strongly matched empirical behavior and psychometrics across all settings (Figures S6A-F). DBS-like stimulation for amplitudes of at least 200 μA regardless of frequency substantially lowered boundary separation (pd>99.05%, median<-0.339, <3.93% in ROPE) and substantially increased drift rate (pd>96.7%, median>1.116, <2.18% in ROPE) (Figure 2E & F). The effect size for both parameters was greatest for 300 μA stimulation at 130 Hz. No setting showed any substantial effect on bias or non-decision time (Figure S6G & H). This modeling suggests that the improvement in conflict resolution is sensitive to the choice of stimulation amplitude and can be resolved within a session.

### Bayesian optimization can robustly identify effective parameters for maximizing cognitive control using DBS-like stimulation

Having confirmed that cognitive control improvements from DBS-like stimulation can be measured acutely, we ran a proof of concept experiment in a new cohort, where stimulation amplitudes were chosen for individual rats by minimizing RTs using Bayesian optimization (Figure 3A, Figure S7). During each Set-Shift trial, rats received stimulation at a specific stimulation amplitude (continuous, but variable amplitude stimulation, in comparison to the intermittent stimulation of prior experiments). After responding, the RT and corresponding amplitude were added to a growing dataset of such pairs. This dataset was used to continuously fit a Gaussian process (GP) model of the individual rat’s RT as a function of stimulation amplitude. An acquisition function balancing exploration and exploitation was computed from this GP and maximized to select the next amplitude to test. 3.5 s into the following inter-trial-interval, the amplitude was updated to the new value before repeating the optimization loop (Figure 3B). In all 6 rats, optimization converged to a non-zero stimulation amplitude that acutely minimized reaction times (Figure 3C & Figure S8). These amplitudes for each rat were comparable to those identified during parallel sessions where settings were selected by brute force space filling (Figure 3D and Figure S9), confirming that the GP identified a reasonable global optimum. Optimization runs converged to their corresponding optimal settings faster than the parallel space filling sessions (Figure 3E). In a follow-up experiment, rats completed Set-Shift while receiving continuous sham, standard 300 μA, or individually optimized stimulation. The settings identified by optimization significantly decreased RTs with a comparable effect to the standard 300 μA parameters (*β* < −52 ms and *P* < 0.014 for both, Figure 3F, Table S11). Neither setting significantly affected accuracy (*P* > 0.05, Figure 3G, Table S12). Together, these results support that Bayesian optimization can rapidly and robustly identify stimulation parameters that acutely improve cognitive control, and that these improvements translate to chronic use.

**Figure 3:**
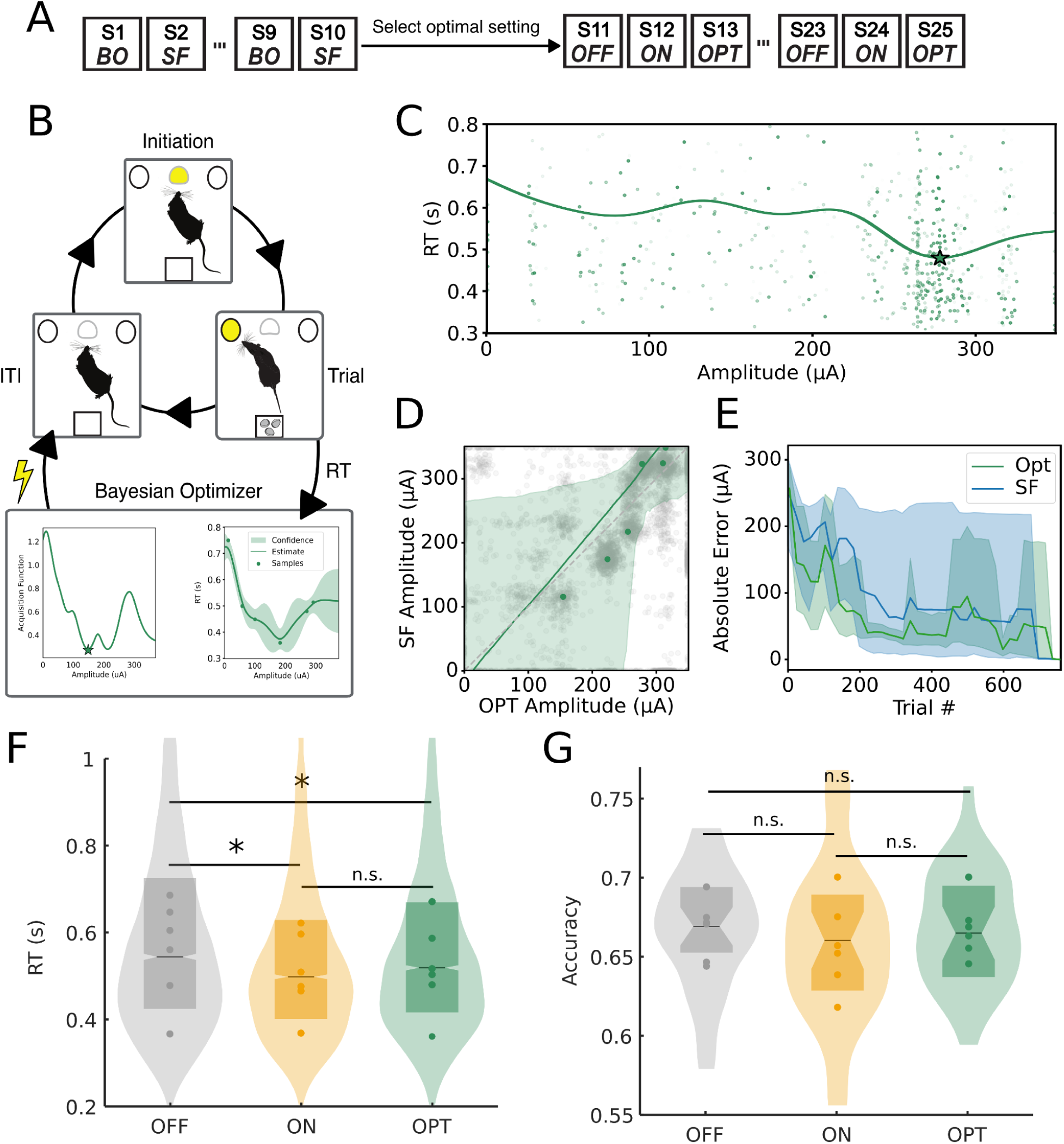
Bayesian optimization can robustly identify effective stimulation amplitudes for enhancing cognitive control. **(A)** Illustration of the sequence of task conditions for the optimization experiment. Rats completed 5 sessions of alternating Bayesian optimization (BO) and brute-force space filling (SF). Following these 10 sessions, each rat’s optimal amplitude was computed from the BO data. In a follow-up experiment, rats completed 5 sessions each of continuous off (OFF), continuous on (ON), or continuous optimized (OPT) stimulation for a total of 15 sessions. Sessions were completed in a predefined repeating sequence of OFF/ON/OPT. **(B)** Illustration of the optimization loop. Rats begin each trial by initiating at the middle port before selecting a side after a period of time (the response time). These response times are used to build a Gaussian process model of the rat’s response time as a function of a corresponding stimulation amplitude. An acquisition function balancing exploration and exploitation is calculated from this model and maximized to select the next amplitude to test. The amplitude is then changed to the new value during the inter-trial-interval before repeating the loop with the new setting. **(C)** Example samples, Gaussian process model, and optimal parameters for an individual rat. The green curve indicates the Gaussian process model. The optimal amplitude is indicated by the green star. Each point in the scatter represents a single trial pairing a response time with the corresponding stimulation amplitude for that trial. The opacity of each point corresponds to the age of the sample relative to the end of the optimization process with more transparent samples being older. **(D)** Correlation between the optimal samples identified by the optimization process as compared to space filling. Each point in gray corresponds to a bootstrapped optimal amplitude from either approach while the points in green are the optimal pairs from the actual data. The line in green represents the line of best fit through all bootstrapped samples while the shaded green region is the 95% confidence interval. **(E)** Convergence of optimization (green) and space filling (blue) over time. At each trial over the optimization process, the current optimal setting is compared to the final optimal setting (absolute error). The shaded region corresponds to the 95% confidence interval over bootstrapped samples. **(F)** Distributions of reaction time as a function of task condition (continuous off, on at 300 μA and 130 Hz, or on at optimal parameters OPT for the individual rat with on target electrodes). Mean values for each rat in each condition are shown by the scattered points. Distributions were computed over all trials. **(G)** Distributions of accuracy as a function of task condition. Distributions were computed over all sessions. *P < 0.05; all P values represent Wald Z tests of model parameters from GLMMs. See supplemental tables for formulae and statistical details.

### DBS-like stimulation shows evidence of trade-offs between cognitively relevant behavioral measures

Lastly, we explored relationships between cognitively relevant behavioral outcomes for different stimulation amplitudes using the space filling data. We considered three outcomes–RT, delay time, and accuracy–and fit GPs for each subject with full hyperparameter optimization. All three outcomes varied smoothly as a function of stimulation amplitude. RT showed a large spread in optimal amplitude across rats although multiple subjects had an optimal setting close to the upper boundary (Figure 4A). The optimal setting for delay time was more consistent across rats and was centered around 200 μA (Figure 4B). The optimal setting for accuracy was more variable but generally required lower amplitudes than for the other two measures (Figure 4C). We plotted Pareto-optimal settings for each rat separately to identify specific tradeoffs between the three outcomes (Figure 4D). In the majority of the rats, raising stimulation amplitudes to the point where RT had a maximal decrease came at the cost of slight reductions in accuracy and increased delay time. However, there were stimulation amplitudes for most rats where a slightly smaller reduction in RT also produced a concurrent reduction in delay time and improvement in accuracy. Together, these results suggest that there is likely a tradeoff between reducing RT and potentially affecting other cognitively relevant outcomes.

**Figure 4:**
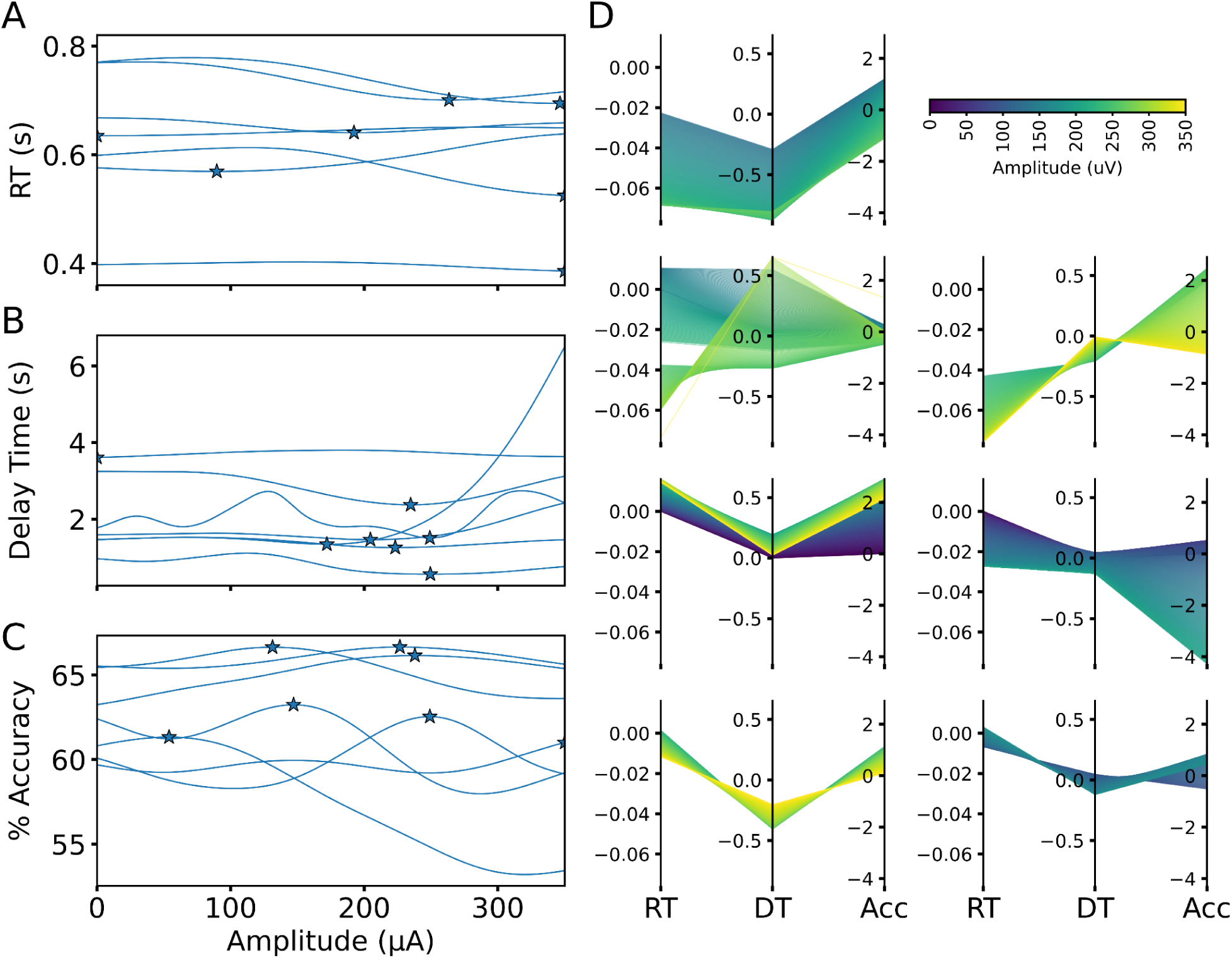
Stimulation at different amplitudes shows evidence of trade-offs between optimizing different cognitive variables. **(A-C)** Best-fitting Gaussian process models of space filling data for all 6 rats for three outcomes of interest: reaction time **(A)**, delay time **(B)**, and accuracy **(C)**. Optimal amplitudes for each rat and outcome are indicated by the blue stars. **(D)** Parallel coordinate plots of pareto optimal settings for all 6 rats. Each pareto optimal setting is indicated by a single polyline connecting the three outcomes and colored by the corresponding stimulation amplitude. All outcomes are normalized relative to the effect at an amplitude of 0 uA for the corresponding rat.

## Discussion

Using our reverse translational model, we showed that black-box optimization can rapidly and robustly identify stimulation parameters that improve a clinically-relevant marker, in this case performance on a cognitive control task. Our approach highlights the power of optimization as a tool for target engagement in psychiatry. Subjective self-reports have historically been inconsistent for effectively tailoring treatment to a patient’s needs [3,4]. We provide a framework for instead approaching this personalization problem from a quantitative perspective with standardized and reproducible methods.

Our Bayesian optimization framework successfully identified stimulation amplitudes that reduced reaction times in all tested animals. Additionally, the acute improvement translated readily to continuous testing. It is worth noting that the improvement from individually optimized parameters was comparable to standard 300 μA stimulation. However, this is not surprising, because we chose 300 μA as our target amplitude in prior work [20] based on months of preliminary experiments manually testing amplitudes for entire sessions with many rats. In contrast, our optimization approach identified comparably effective parameters for individual animals within just 5 days of testing. This more rapid optimization approach would be useful, for instance, to identify parameters that improved a different behavioral or physiologic marker, particularly if those parameters varied across individuals.

In this particular application, we determined that chronically effective parameters could be readily identified based on acute testing. This feature is essential, because it allows for rapid testing of settings rather than requiring hours to days to measure efficacy. With the eventual goal of optimizing multiple parameters including dose, frequency, and location, rigorous testing would not be possible without rapid feedback [22]. However, this approach is not without limitations. In our case, acute and chronic stimulation had comparable effects on RT, delay to initiate a trial, and accuracy. In other applications, like those in a clinical setting, there could be differences that would have to be addressed. This is especially true of the timing for the acute measurements. In our testing, responses to stimulation could be observed in a single trial, but comparable human experiments have shown carry-over between trials, meaning that each stimulation setting would need to be observed over multiple trials [17]. When translating these black-box approaches to clinical trials, ensuring an appropriate balance between the number of measurements and the duration of each measurement will be integral to success.

In our variable parameter testing, 130 Hz and 20 Hz stimulation at 300 μA performed similarly for most metrics. This is surprising, since we found increases in RT and reductions in performance with 20 Hz stimulation in our prior study [20] and low frequency stimulation typically is less effective in movement disorders [36]. The current study tested more than double the animals as in the prior cohort and had a more rigorous testing paradigm which could explain the differences. Additionally, low frequency stimulation is under-explored in psychiatric applications and is beneficial in some cases [49,50].

However, this result could also indicate a difference between acute and chronic measurements dependent on the parameters being tested. Further testing with multi-parameter optimization would be necessary to confirm that acute improvements translate to chronic use across the full-spectrum of stimulation settings.

Our space filling sessions allowed us to assess relationships between multiple outcomes without biasing sampling to optimize a single outcome. We found clear evidence of trade-offs between outcomes, particularly small decreases in accuracy for the largest improvements in reaction time. This matches our earlier finding in the variable parameter experiment where 200 μA stimulation actually improved accuracy with a numerically smaller RT improvement than we observed for 300 μA stimulation. This observation raises a concern for eventual clinical translation: there may be trade-offs between outcomes that could be missed by overly narrow focus on a single measurement [51]. Thus, optimization tools like these will likely be most useful if they can provide clinicians with a set of parameters with a range of improvements in the primary cognitive outcome of interest. Multi-objective optimization approaches may also assist in this endeavor, enabling clinicians to choose parameters with optimal tradeoffs between relevant outcomes [52].

In summary, we show that black-box optimization is a feasible approach for identifying neuromodulation parameters that can engage target circuits to produce desired behavioral and physiologic outcomes. Such tools will allow clinicians to make quantitative decisions during stimulation programming instead of relying on noisy self-reports. The resulting improvements in target engagement should increase the efficacy of neuromodulation therapies, moving closer to the promise of personalized medicine.

## CRediT authorship contribution statement

**Evan M. Dastin-van Rijn:** Writing – original draft, Visualization, Validation, Software, Project administration, Methodology, Investigation, Formal analysis, Data curation, Conceptualization. **Elizabeth M. Sachse:** Writing – review & editing, Validation, Methodology, Investigation. **Michelle Buccini:** Writing – review & editing, Validation, Investigation. **Bradley Angstadt:** Writing – review & editing, Validation, Investigation. **Jonathan Bennek:** Writing – review & editing, Validation, Investigation. **Megan Mensinger:** Writing – review & editing, Validation, Investigation. **Alik S. Widge:** Writing – review & editing, Validation, Supervision, Resources, Project administration, Methodology, Funding acquisition, Conceptualization.

## Data and code availability

Data and code are available on Github at https://github.com/tne-lab/Rat-Optimization-Paper.

