## Supplemental Material for "Real-time Bayesian optimization of deep brain stimulation for personalized cognitive control enhancement"

**The PDF file includes:**

Figures S1 to S9
Tables S1 to S14

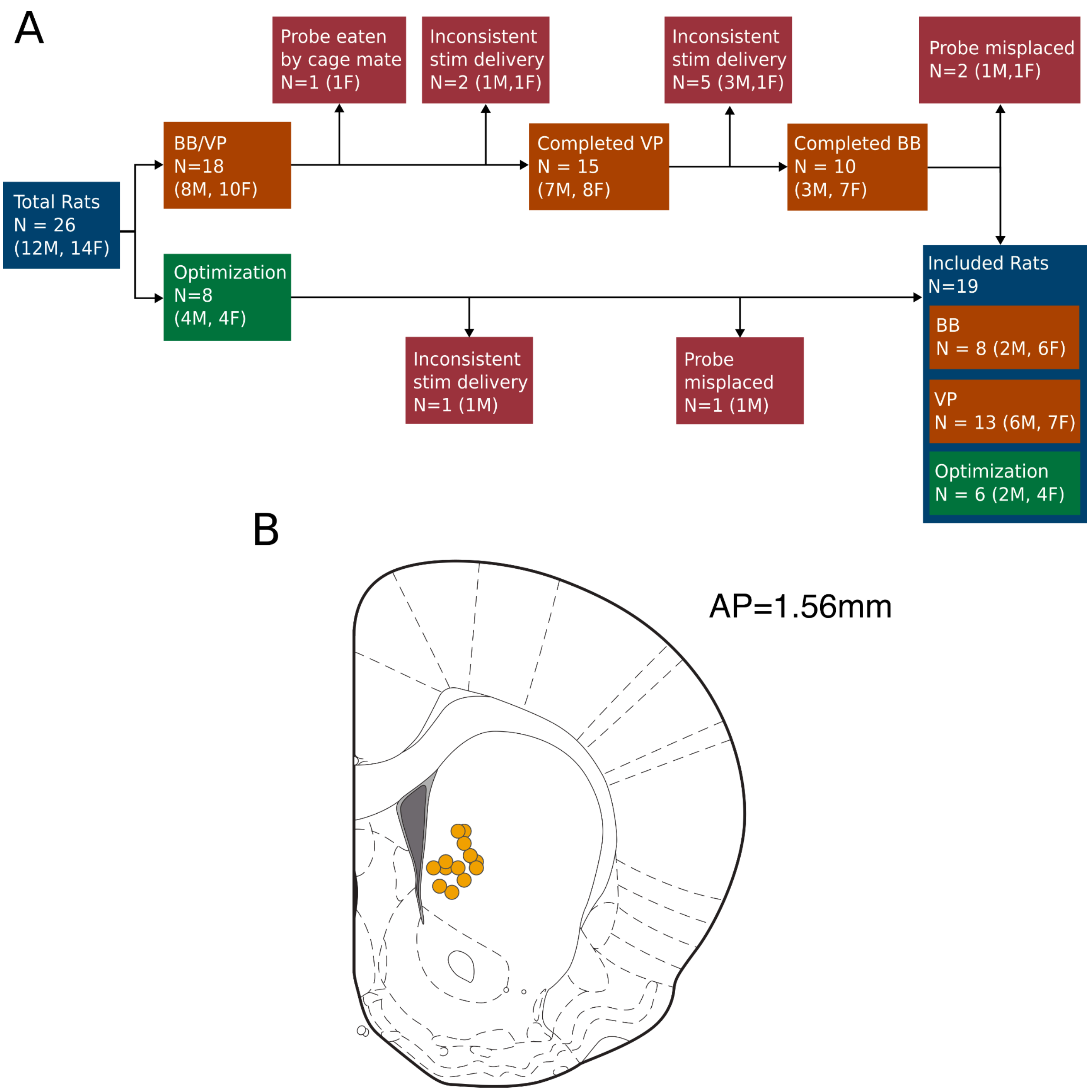


**Figure S1:** Consort diagram and surgical targeting for the BB/VP cohort
**(A)** CONSORT animal flow diagram for the two animal cohorts used in the experiments described in the manuscript: block-based/variable parameter and optimization. Block-based and variable parameter stimulation were tested in the same animals sequentially while optimization was tested in a separate cohort. Any evidence of stimulation cable disconnection or shorting was treated as an exclusion criterion and noted as “inconsistent stim delivery” in the consort diagram. **(B)** Histologically confirmed sites of implant (n = 13 rats total) in the mid-striatum for the rats assigned to the BB/VP cohort.

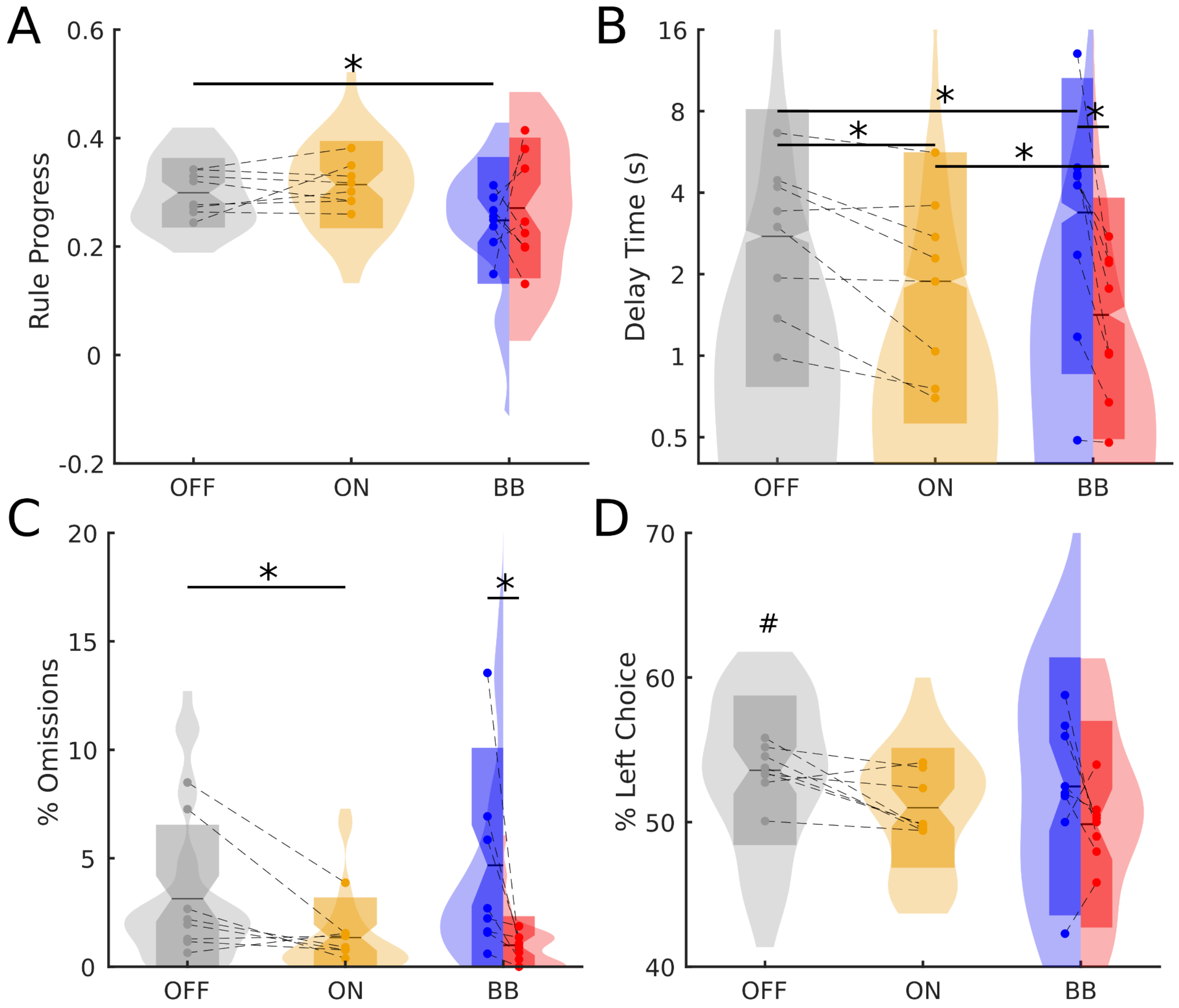


**Figure S2:** Further outcomes of stimulation in the block-based experiment
**(A)** Distributions of the rate at which the rats progressed through the set-shift rules as a function of task condition (rule progress). Distributions were computed over all sessions. **(B)** Distributions of delay time (latency between the middle, cue light turning on and when a rat initiated a trial) as a function of task condition. Mean values for individual rats in each condition are shown by the scattered points and dashed lines link points from the same rats between conditions. Distributions were computed over all trials. **(C)** Distributions of percent of trials with omissions (where rats failed to choose a port within 3 s) as a function of task condition. Distributions were computed over all sessions. **(D)** Distributions of percent of trials where the rat chose the left port as a function of task condition. Distributions were computed over all sessions.
*P < 0.05; all P values represent Wald Z tests of model parameters from GLMMs. See tables S3-S6 for statistical details.

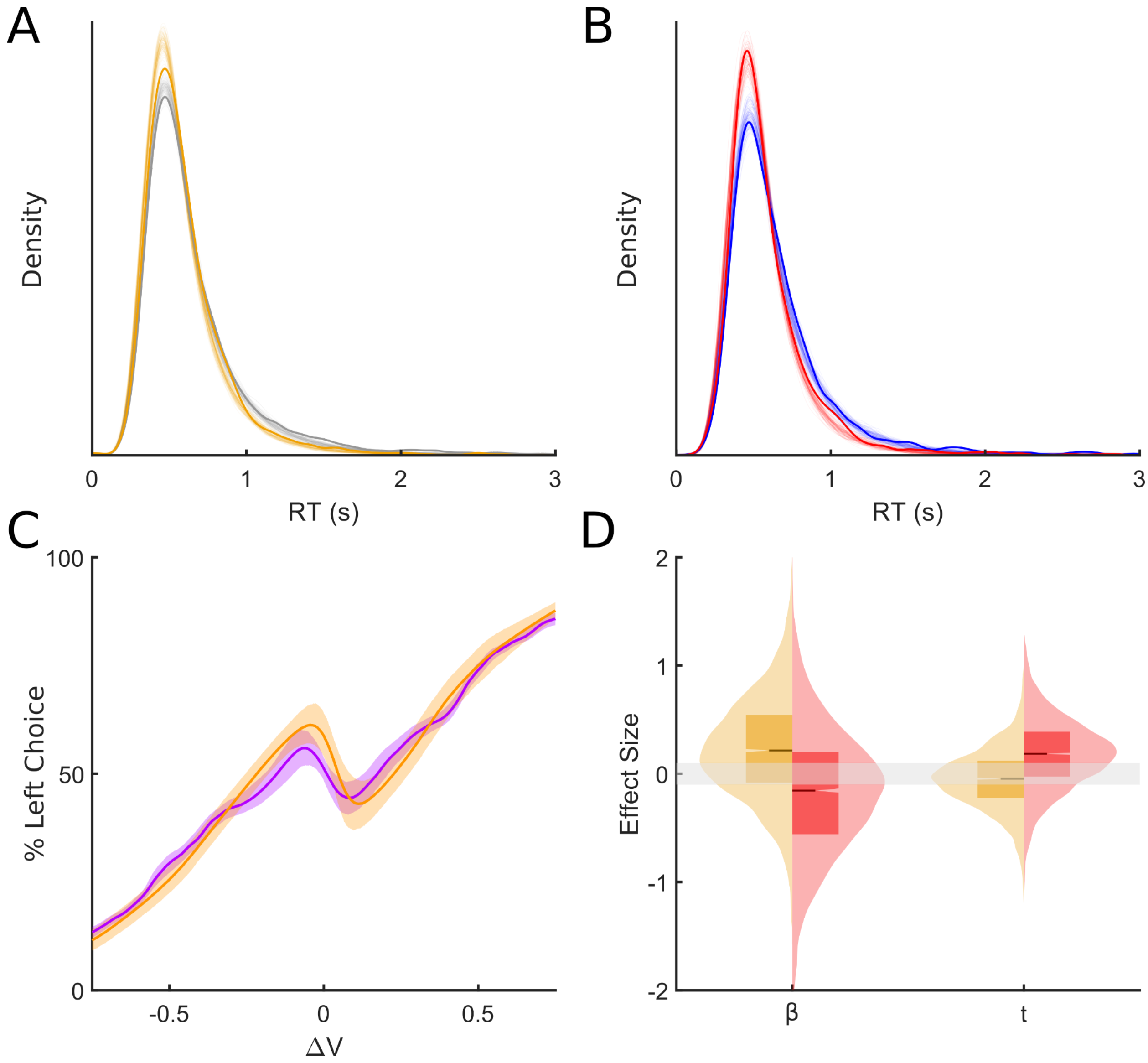


**Figure S3:** Validation of RLDDM fit and additional stimulation effects on model parameters in the block-based experiment

**(A-B)** Distributions of RT from the original data (solid) and model simulations (transparent) for whole-task sham (gray) and whole-task active (yellow) stimulation in **(A)** and block-based sham (blue) and block-based active (red) stimulation in **(B)**. **(C)** Posterior predictive simulations of choice behavior for models (orange) and rats (purple) over 4000 hierarchical posterior draws. Shading shows the 95% highest density interval across animals and trials. Choice behavior was computed as a function of the difference in value between the left and right sides ($\Delta V$). **(D)** Distributions of stimulation effect on bias and non-decision time over 4000 posterior draws. The median of the distribution is indicated by the solid line, the 95% confidence interval by the notch, and the interquartile range by the box. The grey shaded area around zero represents the Region of Practical Equivalence (ROPE) for a null effect (effect size less than 0.1). For both parameters, a sizable portion of the distribution (>10% is in the ROPE) and the distributions don't show consistent directionality (pd is close to 50%).

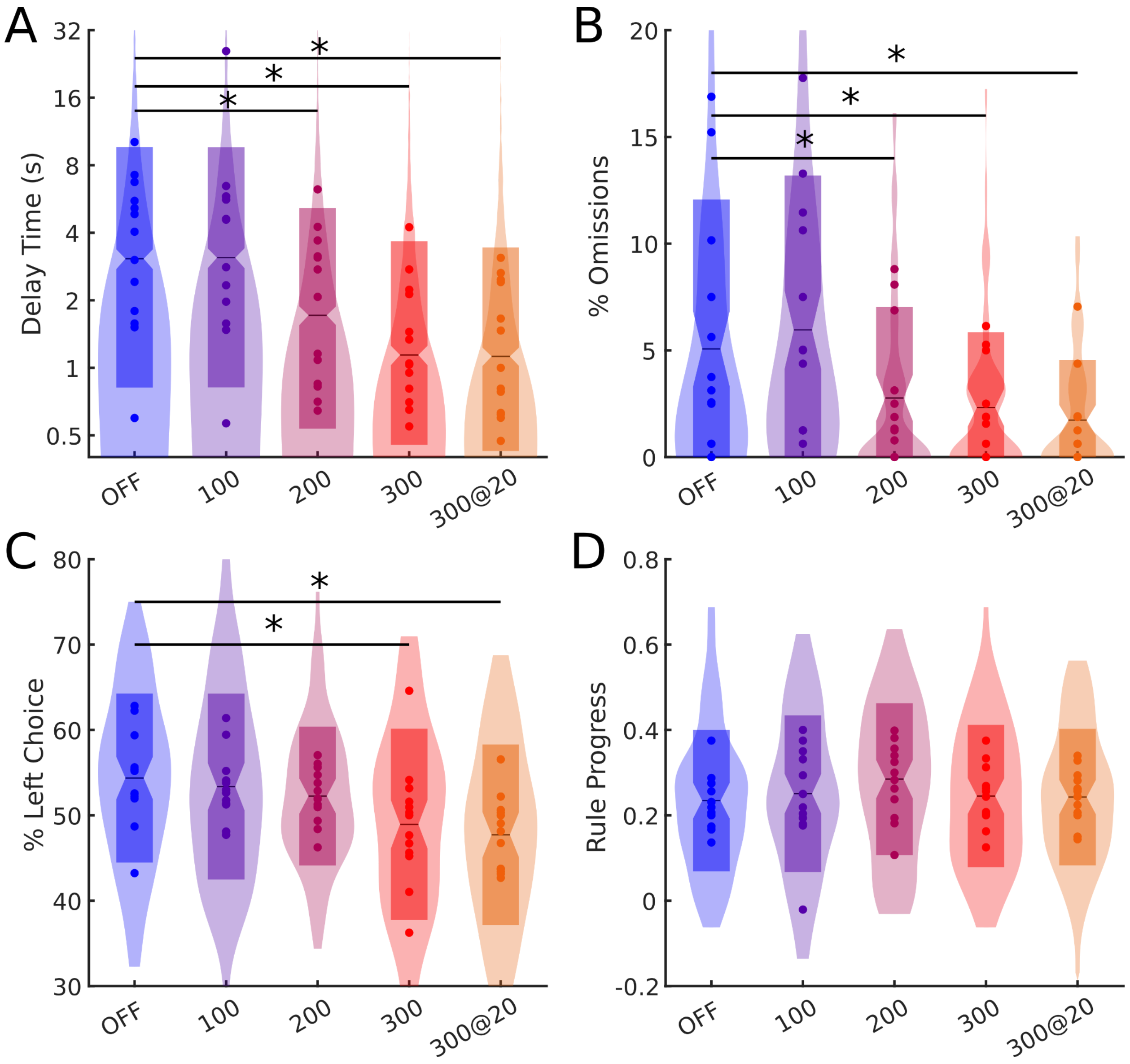


**Figure S4:** Further outcomes of stimulation in the variable parameter experiment

**(A)** Distributions of delay time (latency between the middle, cue light turning on and when a rat initiated a trial) as a function of task condition. Mean values for individual rats in each condition are shown by the scattered points and dashed lines link points from the same rats between conditions. Distributions were computed over all trials. **(B)** Distributions of percent of trials with omissions (where rats failed to choose a port within 3 s) as a function of task condition. Distributions were computed over all sessions. **(C)** Distributions of percent of trials where the rat chose the left port as a function of task condition. Distributions were computed over all sessions. **(D)** Distributions of the rate at which the rats progressed through the set-shift rules as a function of task condition. Distributions were computed over all sessions.

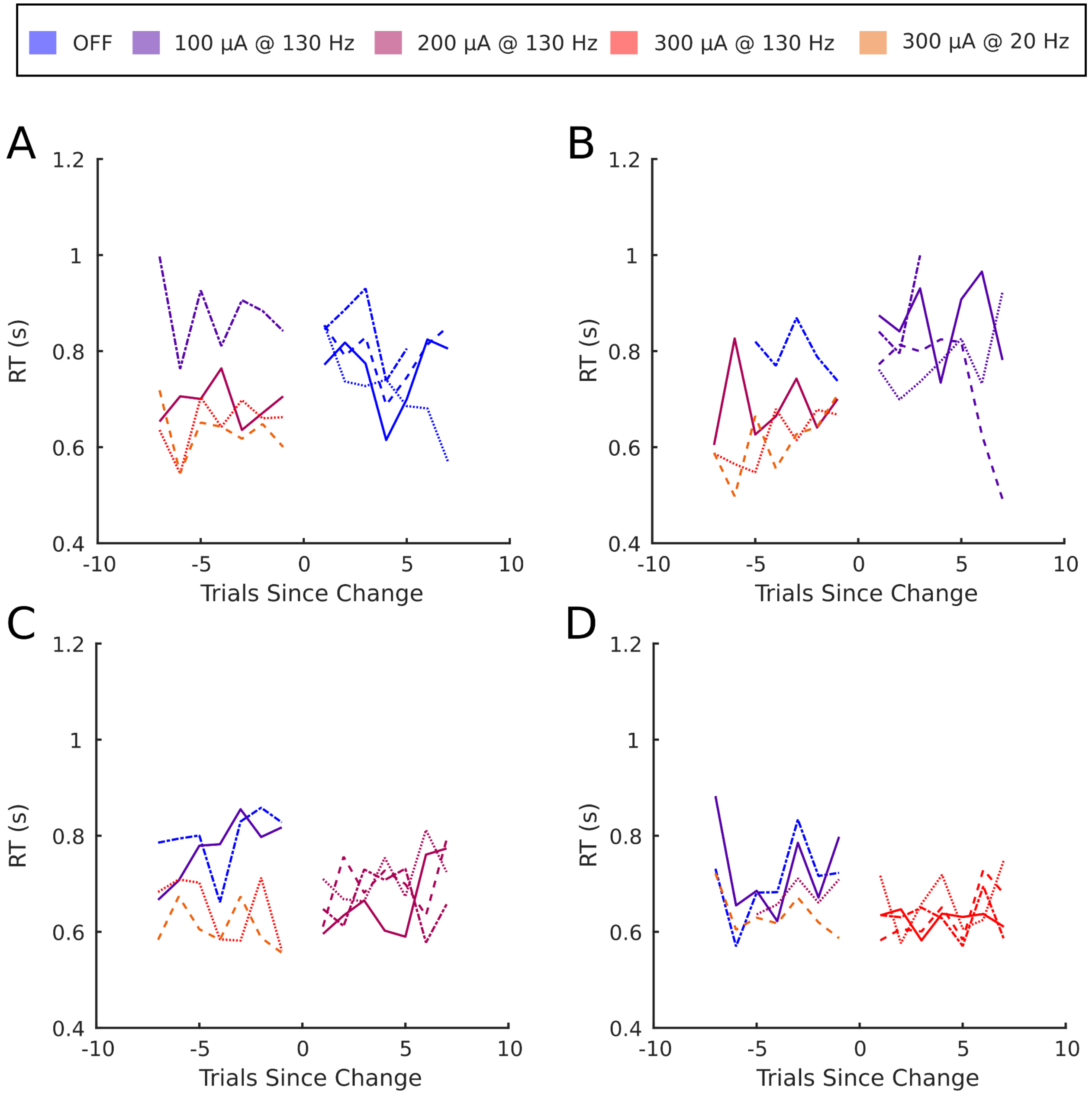


**Figure S5:** Changes in reaction time when updating amplitude during the variable parameter experiment **(A-D)** Change in reaction time (RT) when parameters were updated to the four stimulation conditions not shown in the main text (**A** - 0 µA, **B** - 100 µA, **C** - 200 µA, and **D** - 300 µA at 130 Hz) . Reaction times were averaged separately by the parameters that preceded the corresponding condition. Paired pre and post change averages are indicated by the line-style.

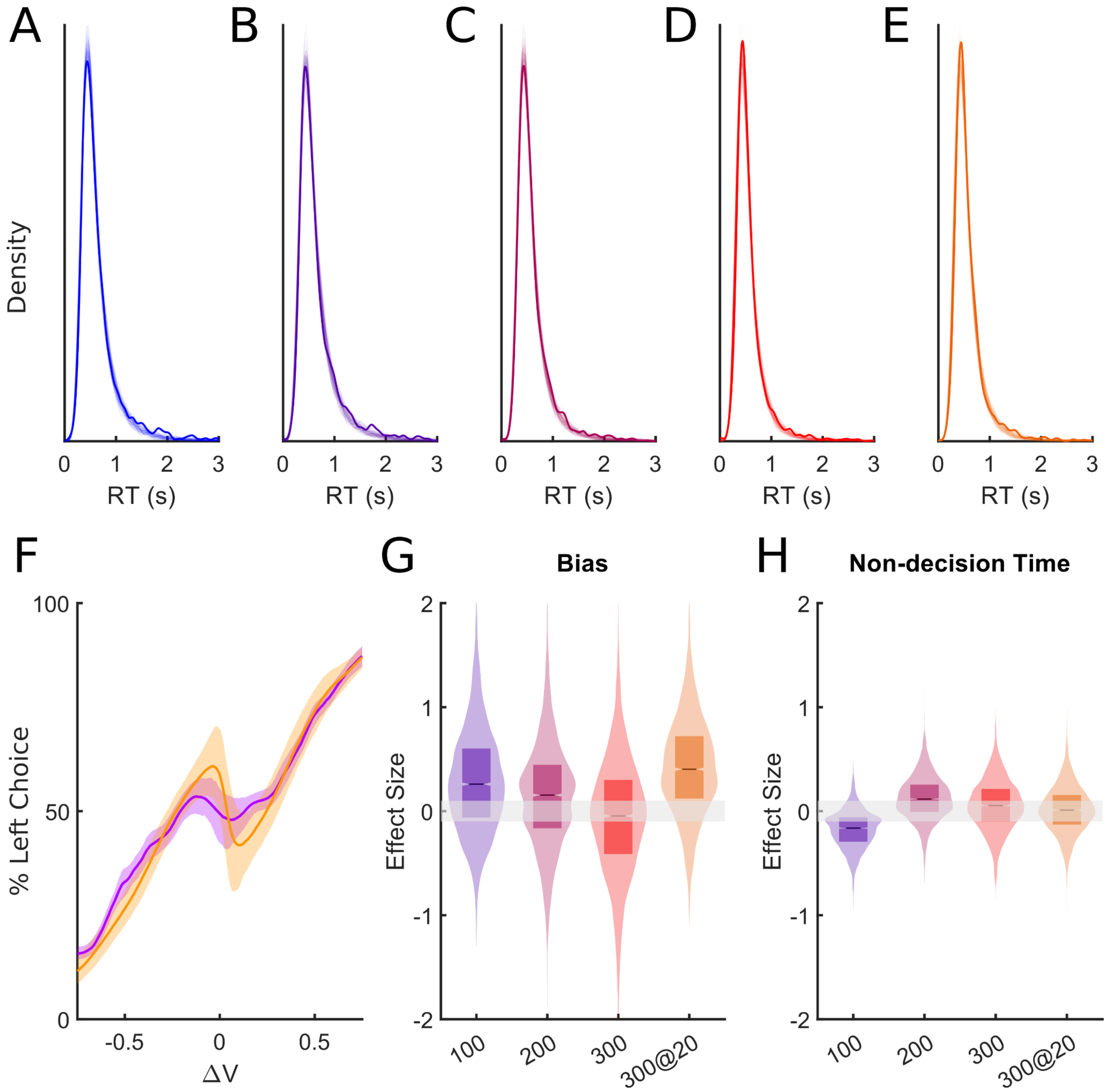


**Figure S6:** Validation of RLDDM fit and additional stimulation effects on model parameters in the variable parameter experiment

**(A-E)** Distributions of RT from the original data (solid) and model simulations (transparent) for all stimulation conditions in the variable parameter experiment (blue=sham, purple=100µA @ 130 Hz, pink=200µA @ 130 Hz, red=300µA @ 130 Hz, and orange=300µA @ 20 Hz). **(F)** Posterior predictive simulations of choice behavior for models (orange) and rats (purple) over 4000 hierarchical posterior draws. Shading shows the 95% highest density interval across animals and trials. Choice behavior was computed as a function of the difference in value between the left and right sides (ΔV). **(G-H)** Distributions of stimulation effect on bias **(G)** and non-decision time **(H)** over 4000 posterior draws. The median of the distribution is indicated by the solid line, the 95% confidence interval by the notch, and the interquartile range by the box. The grey shaded area around zero represents the Region of Practical Equivalence (ROPE) for a null effect (effect size less than 0.1).

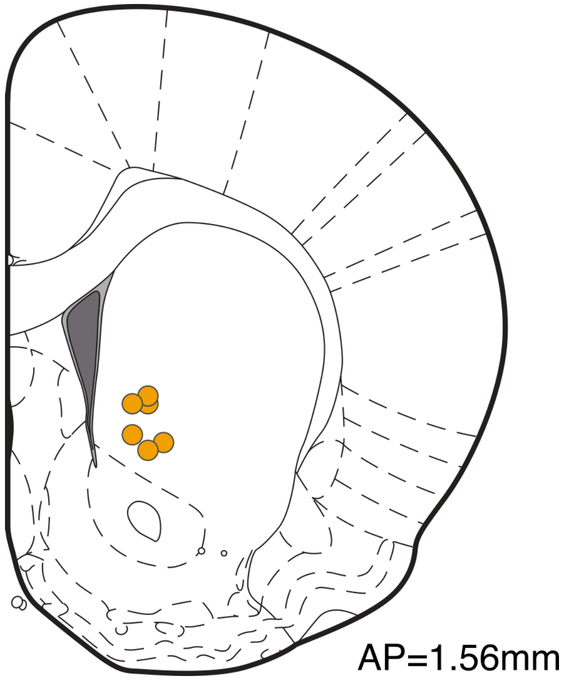


**Figure S7:** Surgical targeting for the optimization cohort
Histologically confirmed sites of implant (n = 6 rats total) in the mid-striatum for the rats assigned to the optimization cohort.

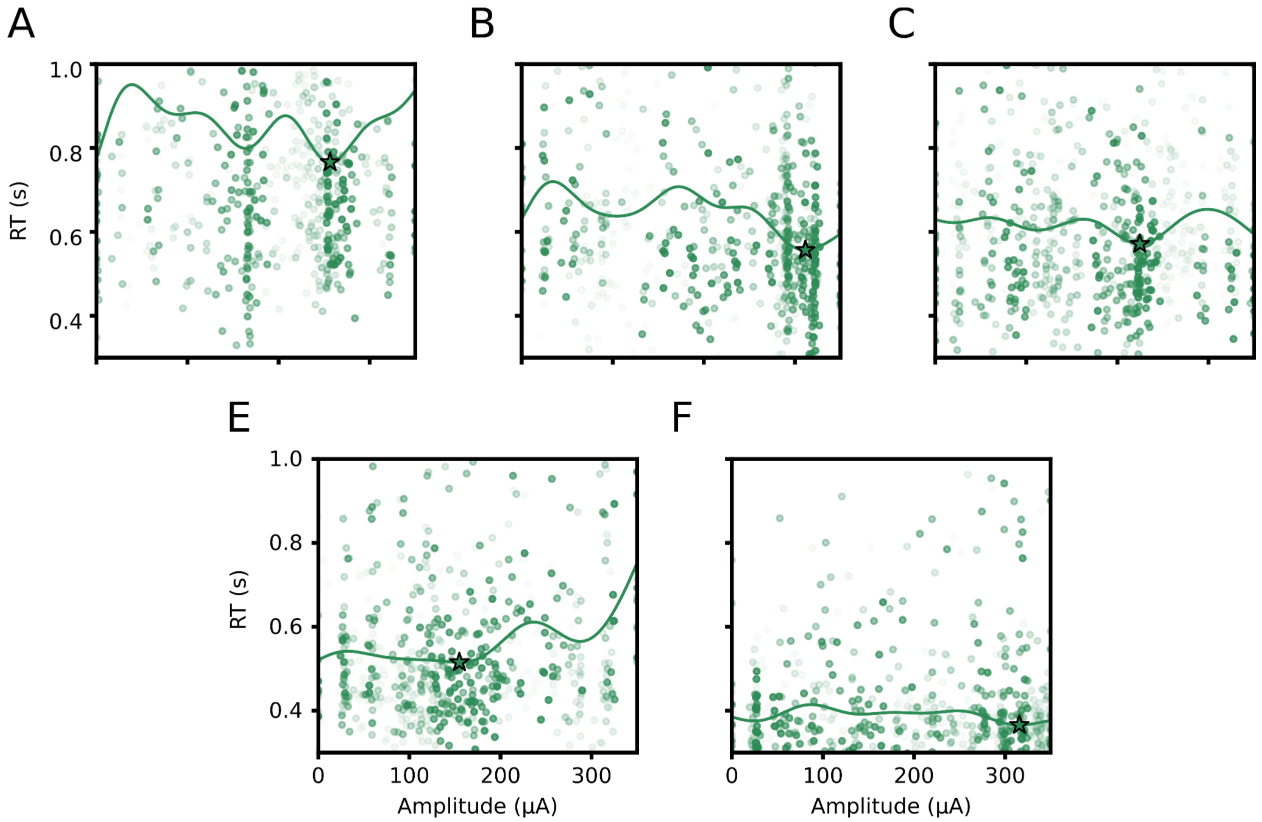


**Figure S8:** Optimization data from additional rats

**(A-F)** Example samples, Gaussian process model, and optimal parameters for the optimization sessions from the 5 rats not included in the main text figure. The green curve indicates the Gaussian process model. The optimal amplitude is indicated by the green star. Each point in the scatter represents a single trial pairing a response time with the corresponding stimulation amplitude for that trial. The opacity of each point corresponds to the age of the sample relative to the end of the optimization process with more transparent samples being older.

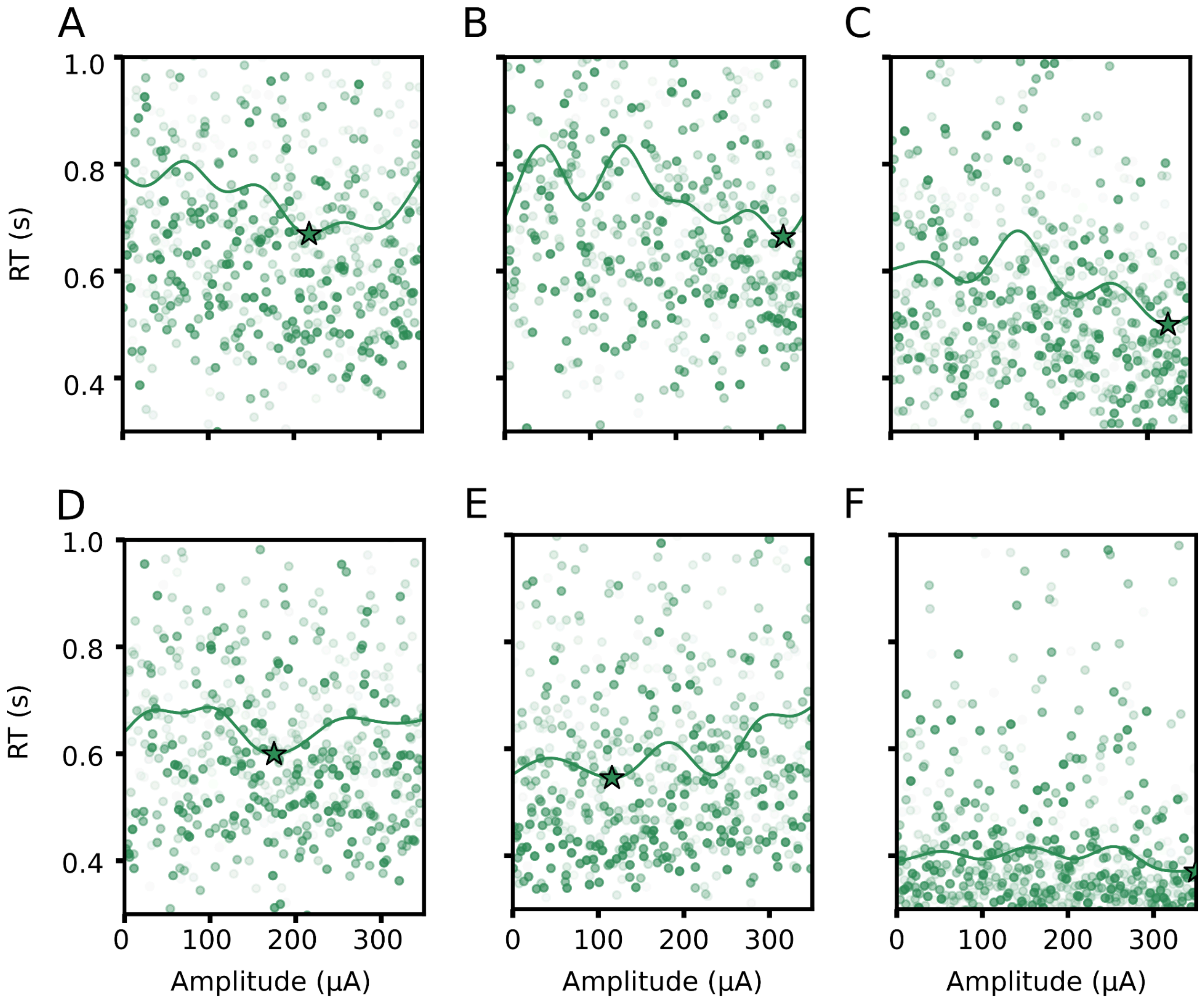


**Figure S9:** Space filling data from all rats

**(A-G)** Example samples, Gaussian process model, and optimal parameters for the space filling sessions from all 6 rats. The green curve indicates the Gaussian process model. The optimal amplitude is indicated by the green star. Each point in the scatter represents a single trial pairing a response time with the corresponding stimulation amplitude for that trial. The opacity of each point corresponds to the age of the sample relative to the end of the optimization process with more transparent samples being older.

**Table S1:** Regression coefficients for reaction time (RT) in seconds in the block-based Set-Shift vs. chronic stimulation experiment, from a generalized linear mixed-effects model, with a gamma distribution and identity link function. Bonferroni adjustments are shown applied to the stimulation coefficients, which were our primary hypothesis and analytic focus. Acute and chronic stimulation both equivalently improved RT.

Formula: RT ~ SessionType + BB_ON + Rule + (1|Subject) + (1|Session) + (1|Protocol)

Fixed effects coefficients:

| **Name** | **Coefficient** | **SE** | **tstat** | **DF** | **p-value** | **Adjusted p** |
|---|---|---|---|---|---|---|
| (Intercept) | 0.667 | 0.040 | 16.58 | 17450 | 2.77e-61 | |
| SessionType_ON | -0.061 | 0.017 | -3.65 | 17450 | 2.60e-4 | 1.56e-3 |
| SessionType_BB | -0.018 | 0.018 | -1.01 | 17450 | 0.31 | 1.00 |
| BB_ON | -0.059 | 0.007 | -7.84 | 17450 | 4.68e-15 | 2.81e-14 |
| Rule_Light | 0.013 | 0.004 | 3.04 | 17450 | 2.34e-3 | |

Random effects coefficients:

| Group | Levels | Estimate |
|---|---|---|
| Subject | 8 | 0.106 |
| Session | 116 | 0.072 |
| Protocol | 5 | 0.018 |

**Table S2:** Regression coefficients for accuracy in the block-based Set-Shift vs. chronic stimulation experiment, from a generalized linear mixed-effects model, with a binomial distribution and logit link function. Bonferroni adjustments are shown applied to the stimulation coefficients, which were our primary hypothesis and analytic focus. No stimulation condition had a significant effect on accuracy.

Formula: Accuracy ~ SessionType + BB_ON + Rule + (1|Subject) + (1|Session) + (1|Protocol)

Fixed effects coefficients:

| **Name** | **Coefficient** | **SE** | **tstat** | **DF** | **p-value** | **Adjusted p** |
|---|---|---|---|---|---|---|
| (Intercept) | 0.359 | 0.059 | 6.31 | 17450 | 2.79e-10 | |
| SessionType_ON | -0.019 | 0.041 | -0.48 | 17450 | 0.63 | 1.00 |
| SessionType_BB | -0.101 | 0.051 | -1.96 | 17450 | 0.05 | 0.30 |
| BB_ON | -0.002 | 0.058 | -0.03 | 17450 | 0.97 | 1.00 |
| Rule_Light | 0.680 | 0.034 | 20.13 | 17450 | 4.24e-89 | |

Random effects coefficients:

| Group | Levels | Estimate |
|---|---|---|
| Subject | 8 | 0.107 |
| Session | 116 | 0.070 |
| Protocol | 5 | 0.063 |

**Table S3:** Regression coefficients for rule progress (rate of task progression) in the block-based Set-Shift vs. chronic stimulation experiment, from a linear mixed-effects model. Bonferroni adjustments are shown applied to the stimulation coefficients, which were our primary hypothesis and analytic focus. No stimulation condition had a significant effect on rule progress.

Formula: Progress ~ SessionType + BB_ON + (1|Subject) + (1|Protocol)

Fixed effects coefficients:

| **Name** | **Coefficient** | **SE** | **tstat** | **DF** | **p-value** | **Adjusted p** |
|---|---|---|---|---|---|---|
| (Intercept) | 0.298 | 0.019 | 15.96 | 147 | 6.73e-34 | |
| SessionType_ON | 0.016 | 0.021 | 0.757 | 147 | 0.45 | 1.00 |
| SessionType_BB | -0.050 | 0.022 | -2.35 | 147 | 0.02 | 0.12 |
| BB_ON | 0.023 | 0.022 | 1.01 | 147 | 0.31 | 1.00 |

Random effects coefficients:

| Group | Levels | Estimate |
|---|---|---|
| Subject | 8 | 0.021 |
| Protocol | 5 | 0.021 |

**Table S4:** Regression coefficients for initiation delay (latency between the cue light and trial initiation) in the block-based Set-Shift vs. chronic stimulation experiment, from a generalized linear mixed-effects model, with a gamma distribution and log link function. Bonferroni adjustments are shown applied to the stimulation coefficients, which were our primary hypothesis and analytic focus. Acute and chronic stimulation both reduced initiation delay.

Formula: DelayTime ~ SessionType + BB_ON + Rule + (1|Subject) + (1|Session) + (1|Protocol)

Fixed effects coefficients:

| **Name** | **Coefficient** | **SE** | **tstat** | **DF** | **p-value** | **Adjusted p** |
|---|---|---|---|---|---|---|
| (Intercept) | 2.288 | 0.227 | 10.06 | 17880 | 9.18e-24 | |
| SessionType_ON | -0.571 | 0.104 | -5.51 | 17880 | 3.58e-8 | 2.15e-7 |
| SessionType_BB | 0.141 | 0.111 | 1.27 | 17880 | 0.20 | 1.00 |
| BB_ON | -1.093 | 0.060 | -18.20 | 17880 | 2.26e-73 | 1.36e-72 |
| Rule_Light | -0.287 | 0.034 | -8.40 | 17880 | 4.69e-17 | |

Random effects coefficients:

| Group | Levels | Estimate |
|---|---|---|
| Subject | 8 | 0.609 |
| Session | 116 | 0.430 |
| Protocol | 5 | 1.815e-5 |

**Table S5:** Regression coefficients for omissions in the block-based Set-Shift vs. chronic stimulation experiment, from a generalized linear mixed-effects model, with a binomial distribution and logit link function. Bonferroni adjustments are shown applied to the stimulation coefficients, which were our primary hypothesis and analytic focus. Acute and chronic stimulation both reduced omissions.

Formula: Omission ~ SessionType + BB_ON + Rule + (1|Subject) + (1|Session) + (1|Protocol)

Fixed effects coefficients:

| **Name** | **Coefficient** | **SE** | **tstat** | **DF** | **p-value** | **Adjusted p** |
|---|---|---|---|---|---|---|
| (Intercept) | -3.708 | 0.169 | -21.96 | 17880 | 1.78e-105 | |
| SessionType_ON | -0.844 | 0.254 | -7.26 | 17880 | 9.10e-4 | 5.46e-3 |
| SessionType_BB | 0.464 | 0.247 | -3.32 | 17880 | 0.06 | 0.36 |
| BB_ON | -1.613 | 0.222 | 1.88 | 17880 | 4.00e-13 | 2.40e-12 |
| Rule_Light | -0.076 | 0.104 | -0.73 | 17880 | 0.47 | |

Random effects coefficients:

| Group | Levels | Estimate |
|---|---|---|
| Subject | 8 | 5.22e-13 |
| Session | 116 | 0.897 |
| Protocol | 5 | 1.064e-6 |

**Table S6:** Regression coefficients for side choice in the block-based Set-Shift vs. chronic stimulation experiment, from a generalized linear mixed-effects model, with a binomial distribution and logit link function. Bonferroni adjustments are shown applied to the stimulation coefficients, which were our primary hypothesis and analytic focus. No stimulation condition had a significant effect on side choice.

Formula: LeftChoice ~ SessionType + BB_ON + Rule + (1|Subject) + (1|Session) + (1|Protocol)

Fixed effects coefficients:

| **Name** | **Coefficient** | **SE** | **tstat** | **DF** | **p-value** | **Adjusted p** |
|---|---|---|---|---|---|---|
| (Intercept) | 0.100 | 0.049 | 2.05 | 17450 | 0.04 | |
| SessionType_ON | -0.103 | 0.043 | -2.43 | 17450 | 0.15 | 0.90 |
| SessionType_BB | -0.049 | 0.052 | -0.94 | 17450 | 0.35 | 1.00 |
| BB_ON | -0.096 | 0.056 | -1.72 | 17450 | 0.09 | 0.54 |
| Rule_Light | 0.114 | 0.031 | 3.63 | 17450 | 2.80e-4 | |

Random effects coefficients:

| Group | Levels | Estimate |
|---|---|---|
| Subject | 8 | 1.27e-8 |
| Session | 116 | 0.099 |
| Protocol | 5 | 0.081 |

**Table S7:** Regression coefficients for reaction time (RT) in seconds in the variable parameter Set-Shift experiment, from a generalized linear mixed-effects model, with a gamma distribution and identity link function. Bonferroni adjustments are shown applied to the stimulation coefficients, which were our primary hypothesis and analytic focus. Stimulation amplitudes of at least 200 µA, regardless of frequency, significantly improved RT.

Formula: RT ~ StimCondition + Rule + (1|Subject) + (1|Session) + (1|Protocol)

Fixed effects coefficients:

| **Name** | **Coefficient** | **SE** | **tstat** | **DF** | **p-value** | **Adjusted p** |
|---|---|---|---|---|---|---|
| (Intercept) | 0.658 | 0.044 | 14.91 | 9544 | 1.10e-49 | |
| StimCondition_100 | 0.002 | 0.009 | 0.19 | 9544 | 0.85 | 1.00 |
| StimCondition_200 | -0.049 | 0.009 | -5.58 | 9544 | 2.54e-8 | 1.52e-7 |
| StimCondition_300 | -0.066 | 0.009 | -7.61 | 9544 | 2.99e-14 | 1.79e-13 |
| StimCondition_300/20 | -0.061 | 0.009 | -6.96 | 9544 | 3.71e-12 | 2.23e-11 |
| Rule_Light | 0.009 | 0.006 | 1.60 | 9544 | 0.11 | |

Random effects coefficients:

| Group | Levels | Estimate |
|---|---|---|
| Subject | 13 | 0.155 |
| Session | 63 | 0.057 |
| Protocol | 5 | 0.005 |

**Table S8:** Regression coefficients for initiation delay (latency between the cue light and trial initiation) in the variable parameter Set-Shift experiment, from a generalized linear mixed-effects model, with a gamma distribution and log link function. Bonferroni adjustments are shown applied to the stimulation coefficients, which were our primary hypothesis and analytic focus. Stimulation amplitudes of at least 200 μA, regardless of frequency, significantly reduced initiation delay.

Formula: DelayTime ~ StimCondition + Rule + (1|Subject) + (1|Session) + (1|Protocol)

Fixed effects coefficients:

| **Name** | **Coefficient** | **SE** | **tstat** | **DF** | **p-value** | **Adjusted p** |
|---|---|---|---|---|---|---|
| (Intercept) | 2.442 | 0.230 | 10.64 | 9895 | 2.70e-26 | |
| StimCondition_100 | -0.011 | 0.071 | -0.16 | 9895 | 0.88 | 1.00 |
| StimCondition_200 | -0.715 | 0.071 | -10.06 | 9895 | 1.11e-23 | 6.66e-23 |
| StimCondition_300 | -1.101 | 0.071 | -15.51 | 9895 | 1.22e-53 | 7.32e-53 |
| StimCondition_300/20 | -1.125 | 0.071 | -15.81 | 9895 | 1.21e-55 | 7.26e-55 |
| Rule_Light | 0.135 | 0.047 | -2.88 | 9895 | 3.96e-3 | |

Random effects coefficients:

| Group | Levels | Estimate |
|---|---|---|
| Subject | 13 | 0.790 |
| Session | 63 | 0.240 |
| Protocol | 5 | 0.062 |

**Table S9:** Regression coefficients for omissions in the variable parameter Set-Shift experiment, from a generalized linear mixed-effects model, with a binomial distribution and logit link function. Bonferroni adjustments are shown applied to the stimulation coefficients, which were our primary hypothesis and analytic focus. Stimulation amplitudes of at least 200 μA, regardless of frequency, significantly reduced omissions.

Formula: Omission ~ StimCondition + Rule + (1|Subject) + (1|Session) + (1|Protocol)

Fixed effects coefficients:

| **Name** | **Coefficient** | **SE** | **tstat** | **DF** | **p-value** | **Adjusted p** |
|---|---|---|---|---|---|---|
| (Intercept) | -3.25 | 0.303 | -10.70 | 9895 | 1.39e-26 | |
| StimCondition_100 | 0.186 | 0.147 | 1.27 | 9895 | 0.205 | 1.00 |
| StimCondition_200 | -0.669 | 0.178 | -3.76 | 9895 | 1.72e-4 | 1.03e-3 |
| StimCondition_300 | -0.838 | 0.187 | -4.48 | 9895 | 7.71e-6 | 4.63e-5 |
| StimCondition_300/20 | -1.16 | 0.206 | -5.61 | 9895 | 2.05e-8 | 1.23e-7 |
| Rule_Light | 0.603 | 0.045 | -2.72 | 9895 | 6.46e-3 | |

Random effects coefficients:

| Group | Levels | Estimate |
|---|---|---|
| Subject | 13 | 0.971 |
| Session | 63 | 0.427 |
| Protocol | 5 | 1.437e-16 |

**Table S10:** Regression coefficients for accuracy in the variable parameter Set-Shift experiment, from a generalized linear mixed-effects model, with a binomial distribution and logit link function. Bonferroni adjustments are shown applied to the stimulation coefficients, which were our primary hypothesis and analytic focus. Stimulation at 200 µA significantly improved accuracy.

Formula: Accuracy ~ StimCondition + Rule + (1|Subject) + (1|Session) + (1|Protocol)

Fixed effects coefficients:

| **Name** | **Coefficient** | **SE** | **tstat** | **DF** | **p-value** | **Adjusted p** |
|---|---|---|---|---|---|---|
| (Intercept) | 0.226 | 0.071 | 3.16 | 9552 | 1.59e-3 | |
| StimCondition_100 | 0.113 | 0.069 | 1.65 | 9552 | 0.10 | 0.60 |
| StimCondition_200 | 0.192 | 0.069 | 2.79 | 9552 | 5.21e-3 | 0.031 |
| StimCondition_300 | 0.013 | 0.068 | 0.19 | 9552 | 0.85 | 1.00 |
| StimCondition_300/20 | 0.033 | 0.068 | 0.49 | 9552 | 0.62 | 1.00 |
| Rule_Light | 0.603 | 0.045 | 13.36 | 9552 | 2.54e-40 | |

Random effects coefficients:

| Group | Levels | Estimate |
|---|---|---|
| Subject | 13 | 0.106 |
| Session | 63 | 0.085 |
| Protocol | 5 | 0.084 |

**Table S11:** Regression coefficients for side choice in the variable parameter Set-Shift experiment, from a generalized linear mixed-effects model, with a binomial distribution and logit link function. Bonferroni adjustments are shown applied to the stimulation coefficients, which were our primary hypothesis and analytic focus. Stimulation at 300 µA, regardless of frequency, significantly affected side choice.

Formula: LeftChoice ~ StimCondition + Rule + (1|Subject) + (1|Session) + (1|Protocol)

Fixed effects coefficients:

| **Name** | **Coefficient** | **SE** | **tstat** | **DF** | **p-value** | **Adjusted p** |
|---|---|---|---|---|---|---|
| (Intercept) | 0.108 | 0.056 | 1.92 | 9552 | 0.05 | |
| StimCondition_100 | -0.024 | 0.066 | -0.36 | 9552 | 0.72 | 1.00 |
| StimCondition_200 | -0.072 | 0.065 | -1.10 | 9552 | 0.27 | 1.00 |
| StimCondition_300 | -0.203 | 0.065 | -3.13 | 9552 | 1.77e-3 | 0.01 |
| StimCondition_300/20 | -0.245 | 0.065 | -3.76 | 9552 | 1.70e-4 | 1.02e-3 |
| Rule_Light | 0.124 | 0.042 | 2.96 | 9552 | 3.12e-3 | |

Random effects coefficients:

| Group | Levels | Estimate |
|---|---|---|
| Subject | 13 | 0.064 |
| Session | 63 | 0.015 |
| Protocol | 5 | 0.039 |

**Table S12:** Regression coefficients for rule progress (rate of task progression) in the variable parameter Set-Shift experiment, from a linear mixed-effects mode. Bonferroni adjustments are shown applied to the stimulation coefficients, which were our primary hypothesis and analytic focus. No stimulation condition significantly affected rule progress.

Formula: Progress ~ StimCondition + (1|Subject) + (1|Protocol)

Fixed effects coefficients:

| **Name** | **Coefficient** | **SE** | **tstat** | **DF** | **p-value** | **Adjusted p** |
|---|---|---|---|---|---|---|
| (Intercept) | 0.234 | 0.022 | 10.79 | 310 | 2.89e-23 | |
| StimCondition_100 | 0.016 | 0.030 | 0.55 | 310 | 0.58 | 1.00 |
| StimCondition_200 | 0.050 | 0.030 | 1.68 | 310 | 0.09 | 0.54 |
| StimCondition_300 | 0.011 | 0.030 | 0.36 | 310 | 0.72 | 1.00 |
| StimCondition_300/20 | 0.008 | 0.030 | 0.28 | 310 | 0.78 | 1.00 |

Random effects coefficients:

| Group | Levels | Estimate |
|---|---|---|
| Subject | 13 | 0.015 |
| Protocol | 5 | 1.26e-5 |

**Table S13:** Regression coefficients for reaction time (RT) in seconds in the Set-Shift optimization-follow-up experiment, from a generalized linear mixed-effects model, with a gamma distribution and identity link function. Bonferroni adjustments are shown applied to the stimulation coefficients, which were our primary hypothesis and analytic focus. Optimized and standard amplitudes both significantly improved RT.

Formula: RT ~ SessionType + Rule + (1|Subject) + (1|Session) + (1|Protocol)

Fixed effects coefficients:

| **Name** | **Coefficient** | **SE** | **tstat** | **DF** | **p-value** | **Adjusted p** |
|---|---|---|---|---|---|---|
| (Intercept) | 0.608 | 0.045 | 13.42 | 13124 | 8.63e-41 | |
| SessionType_STD | -0.068 | 0.019 | -3.55 | 13124 | 3.86e-4 | 7.72e-4 |
| SessionType_OPT | -0.052 | 0.019 | -2.71 | 13124 | 6.75e-3 | 0.014 |
| Rule_Light | 0.013 | 0.004 | 3.14 | 13124 | 1.72e-3 | |

Random effects coefficients:

| Group | Levels | Estimate |
|---|---|---|
| Subject | 6 | 0.106 |
| Session | 86 | 0.069 |
| Protocol | 5 | 3.159e-6 |

**Table S14:** Regression coefficients for accuracy in the Set-Shift optimization-follow-up experiment, from a generalized linear mixed-effects model, with a binomial distribution and logit link function. Bonferroni adjustments are shown applied to the stimulation coefficients, which were our primary hypothesis and analytic focus. Neither stimulation condition had a significant effect on accuracy

Formula: Accuracy ~ SessionType + Rule + (1|Subject) + (1|Session) + (1|Protocol)

Fixed effects coefficients:

| **Name** | **Coefficient** | **SE** | **tstat** | **DF** | **p-value** | **Adjusted p** |
|---|---|---|---|---|---|---|
| (Intercept) | 0.442 | 0.042 | 10.45 | 13124 | 1.77e-25 | |
| SessionType_STD | -0.029 | 0.046 | -0.64 | 13124 | 0.52 | 1.00 |
| SessionType_OPT | 0.020 | 0.046 | 0.45 | 13124 | 0.66 | 1.00 |
| Rule_Light | 0.560 | 0.038 | 14.58 | 13124 | 9.01e-48 | |

Random effects coefficients:

| Group | Levels | Estimate |
|---|---|---|
| Subject | 6 | 1.37e-8 |
| Session | 86 | 3.617e-6 |
| Protocol | 5 | 0.051 |
